# A Computational Approach Reveals the Ability of Amyloids to Sequester RNA: the Alpha Synuclein Case

**DOI:** 10.1101/2022.09.20.508776

**Authors:** Jakob Rupert, Michele Monti, Elsa Zacco, Gian Gaetano Tartaglia

## Abstract

Nucleic acids can act as potent modulators of protein aggregation, and RNA is able to either hinder or facilitate protein assembly depending on the molecular context. Here we used a computational approach to characterize the physico-chemical properties of regions involved in amyloid aggregation. In different experimental datasets we observed that, while the core is hydrophobic and highly ordered, external regions, more disordered, display a distinct tendency to interact with nucleic acids.

To validate our predictions, we performed aggregation assays with α-synuclein (aS140), a non-nucleic acid binding amyloidogenic protein, and a mutant truncated at the acidic C-terminus (aS103) that is predicted to sequester RNA. For both aS140 and aS103 we observed acceleration of the aggregation upon RNA addition with a significantly stronger effect for aS103. Due to the favorable electrostatics, we observed enhanced nucleic-acid sequestration ability for aS103 that entrapped a larger amount of RNA.

Overall, our research suggests that RNA sequestration is a rather common phenomenon linked to protein aggregation and constitutes a gain-of-function mechanism to be further investigated.

**STATEMENT OF SIGNIFICANCE:** Our study indicates that aggregation confers RNA-binding ability to non-RNA-binding proteins such as alpha synuclein. The sequestration of RNA upon protein aggregation might alter RNA homeostasis and impact multiple biochemical cascades.

## INTRODUCTION

The order and structural robustness of amyloids render them a powerful platform for attracting other cellular factors and macromolecules^1,2^. Nucleic acids are embroiled in the modulation of protein phase transitions^3,4^, as recent experiments show that they can be isolated from both physiological condensates and pathological inclusions^5,6^. RNA in particular might affect aggregate accumulation rate and nature^7^. In support of this observation, numerous proteins found aggregated in neurological disorders are RNA-binding and form ribonucleoprotein assemblies^4^, and their ability to aggregate appears to be at least in part determined by the type and amount of RNA found in their proximity^1^. Yet, only a few studies have focused on understanding the ability of proteins to interact with nucleic acids after the transition towards amyloid species has taken place, or whether such transition might confer the ability to attract nucleic acids to proteins not classified as RNA/DNA-binding^8^.

Many canonical RNA-binding proteins (RBPs) have a modular structure with one or more RNA-binding domains (RBDs) and at least one prion-like domain able to drive phase transitions^9^. Prion regions represent the core of the amyloid fibril structure upon aggregation^10,11^ and the RBDs, usually found at least partially folded in the aggregated species, are external to the core and might retain RNA-binding ability^12^. This is the case of hnRNPDL-2, for which a cryo-EM structure of the aggregate shows two RBDs in a disordered, solvent-exposed portion of fibrils outside of the amyloid core^12^-the “fuzzy coat”. hnRNPDL-2 fibrils have a comparable affinity towards oligonucleotides both in the monomeric and the aggregated form^12^, confirming that canonical RBP aggregates can retain their functionality even upon partial refolding into different conformations.

For non-canonical RBPs and for proteins that do not display RNA binding activity the functional consequences of amyloid formation are less clear. In the case of microtubule-associated protein tau, it has been shown that fibrils can sequester RNA both in cell^6^ and *in vitro*^13^, implying the protein conformational changes upon aggregation can promote RNA binding. This observation is further strengthened by the fact that tau fibrils, grown *in vitro* and without any additional cofactors, can sequester RNA^14^. Tau is not known as a canonical RBP, however it has been shown that RNA co-localises with this protein^15^ and promotes its aggregation^16^ into morphologically distinct fibril strains^17^. A recent cryo-EM structure has shown that RNA can also trigger the formation of a previously undetected tau aggregate morphology with a central core spanning residues E391 to A426^13^. In previously solved tau fibril structures, this specific segment has been reported to be part of the fuzzy coat, implying that RNA can actively change the fibril morphology. Moreover, electron density showed an unknown density that could correspond to the actual RNA molecules being sequestered in the fibrils.

In this work, we use a computational approach to investigate nucleic acid sequestration in protein aggregates by considering the physico-chemical properties of their amyloid cores, hidden within the fibers and occupied in cross-β bonds, and their exterior, available to establish novel interactions and exert new functions. We validated our model on alpha synuclein (aS) that represents a particularly suitable case, since its phase transition is abundantly studied and does not bind RNA in the monomeric state under physiological conditions. aS is an intrinsically disordered protein involved in synaptic vesicle trafficking and machinery assembly, mitochondrial homeostasis and DNA repair^18–20^. It has been identified as the primary component of Lewy bodies^21^, pathological aggregates found in certain neurodegenerative conditions such as Parkinson’s Disease (PD) and Multiple System Atrophy (MSA)^21–23^. aS has a modular organization, with a basic N-terminus (residues 1-95), containing the amyloid central region (NAC, residues 61-95)^24^, and an acidic, intrinsically disordered C-terminus (residues 96-140), shown to negatively affect fibrillation and oligomerization^25–27^. The C-terminal domain can undergo several post-translational modifications such as truncations at various residues^28,29^, which occur naturally in the cell, presumably as incomplete protein degradation products. However, certain disease-related C-terminal truncations also boost the protein fibrillation rates and result in diverse aggregate morphologies, leading to an increase in cellular toxicity^26,30–32^. In the full length protein, long-range electrostatic interactions between the N-terminus and C-terminus influence the partially folded intermediary states of the protein and are highly affected by changes in pH and ionic strength^27,33–35^. These factors, as well as the presence of various charged polymers, strongly affect the aggregation rates, highlighting the role of electrostatic charge in aS misfolding^36–39^. In the absence of the acidic C-terminus, such in the case of disease-related truncations, it is possible that polyanions such as RNA could have an even more significant impact on aS aggregation.

Based on our calculations, we hypothesize that amyloid-forming proteins, such as aS, can acquire the ability to sequester RNA upon aggregation. We propose that RNA binding is an acquired ability of aS aggregates and that this is the crucial contribution recapitulating the experimental trend in aggregation rate modulation. More in general, our work indicates that misfolding upon aggregation has the potential to vary the RNA-binding ability of any given protein, potentially enabling a non-RNA-binding protein to acquire the ability to bind and sequester RNA in its aggregated form.

## RESULTS

### Calculations recapitulate experimental results on the ability of tau fibrils to attract RNA

We started by assessing the robustness of our Zygregator-based calculations on the known case of tau. A cryo-EM structure of tau fibrils (PDB 7SP1), grown in the presence of total RNA, shows that RNA is trapped in a pocket on the external region of the aggregate^13^. The binding occurs through cation-π interactions between an arginine residue and the RNA phosphate backbone, as well as polar contacts between the RNA bases and the histidine residue of tau^13^. The 36-amino acid region of the C-terminal disordered domain (residues 391 - 426) is identified in the cryo-EM study as the fibril core^13^ and another work indicates that the third microtubule-binding repeat (residues 306 – 335)^59,60^ contributes to tau aggregation. Using the Zyggregator algorithm (**Materials and Methods**), which predicts aggregation-prone elements in proteins^41^, we are able to successfully identify both these regions (**Fig. 1A**).

**Figure 1.**
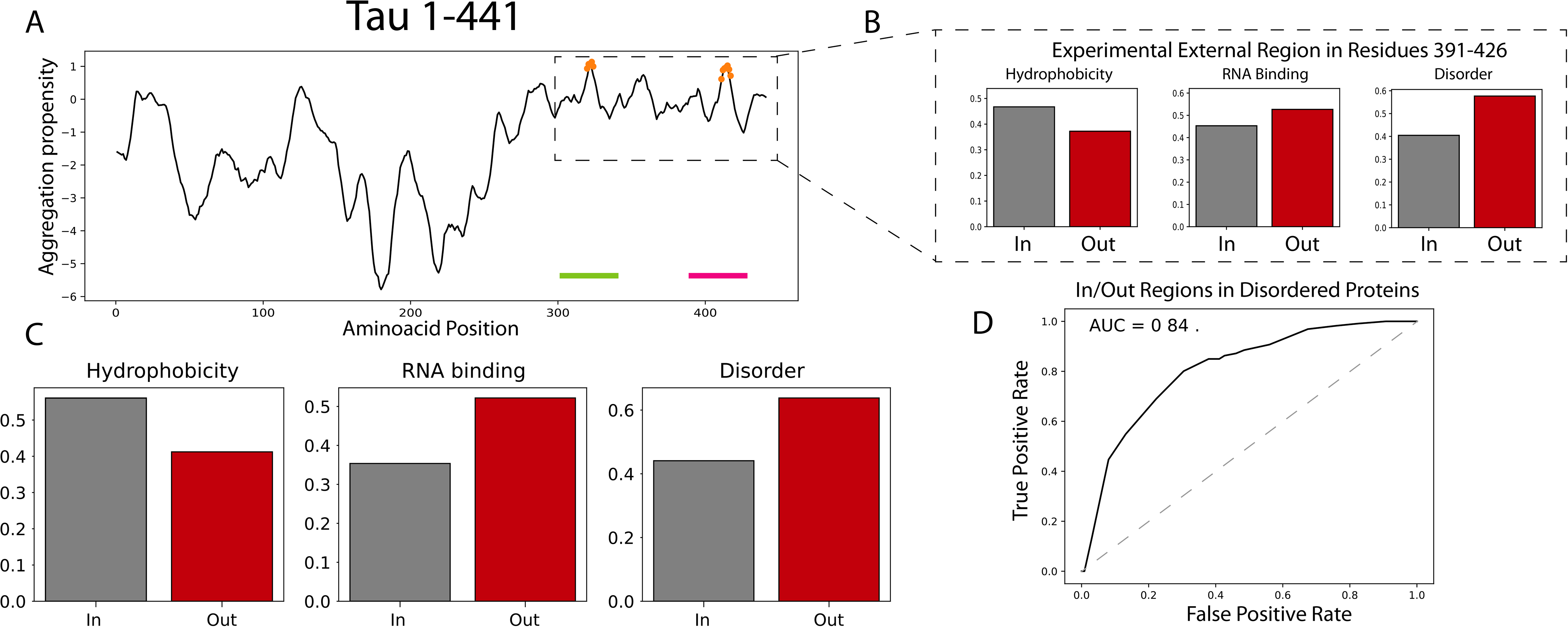
Tau aggregation profile and prediction of physico-chemical properties. **A)** Aggregation propensity profile of tau, with the predictions of the filament internal part in orange, while green and red bars indicate the regions found to be internal experimentally. **B)** Physico-chemical properties of residues 391 - 441, with the internal and external properties of the region marked in black (in) and red (out), respectively. **C)** Hydrophobicity, disorder, and RNA binding propensity of the entire sequence. **D)** ROC curve computed to determine the optimal Zyggregator threshold for finding internal and external residues based on experimental data from various proteins. The optimal Zyggregator threshold is in the range 0.5-1.5.

More in detail, the cryo-EM structure of aggregated tau reveals that residues R406 and H407 on the tau fibril surface directly interact with sequestered RNA and that residues E391 - S396, V398, S400, D402, S404, D421, P423, I425, A426 are exposed to the solvent. In agreement with this finding, we predict that the surface regions identified experimentally are less hydrophobic, more disordered and have an increased RNA-binding ability (**Fig. 1B; Materials and Methods**).

We note that residues 175-220 and 230-260, corresponding to the proline-rich region and first microtubule-binding repeat, have been reported to be directly involved in interactions with RNA^61^ and identified as such by our approach (**Fig. 1C**). Although these disordered regions are not captured by cryo-EM, as not in the stable amyloid core, it cannot be excluded that they play a role in sequestering RNA in the course of aggregation, as also remarked by the authors of the study^13^.

Thus, this initial result indicates that regions outside the core of tau amyloid fibrils have the potential to interact with RNA and we are able to identify them in great detail using our computational approaches. This finding also suggests that other aggregating proteins might behave as tau. For a number of other disordered proteins, including calcitonin^42,43^, amylin^44^, glucagon^45^, Aβ42^46,47^, α-synuclein^48–50^, hnRNPDL-2^12^ and tau^13^, Zyggregator shows excellent performances in identifying the amyloid core of the aggregates, with an optimal Zyggregator threshold in the range 0.5-1.5 (**Fig. 1D**).

### Internal and external regions of amyloids have different biophysical properties

To further investigate the general trend that regions able to bind RNA are in the external parts of amyloids, we retrieved a dataset of amyloids available from a recent publication^62^ and analyzed their physico-chemical properties (**Supplementary Table 1**) using the Zyggregator algorithm to identify regions forming the amyloid core (**Materials and Methods)**.

As expected, our calculations show that the internal part of an amyloid is more hydrophobic than the external one **(Fig. 2A**). There is a significant shift of the RNA-binding propensity and structural disorder in favor of the external part (**Fig. 2B)** while the electrostatic charge distribution does not differ substantially (**Supplementary Fig. 1A,B**). Notably, the trend is observed with different physico-chemical propensity predictors (**Supplementary Fig. 2**) Given the overall negative charge of the phosphate backbone of RNA and no significant difference in the aggregate charge distribution, we find strong RNA-binding propensity exclusively for positively charged protein regions.

**Figure 2.**
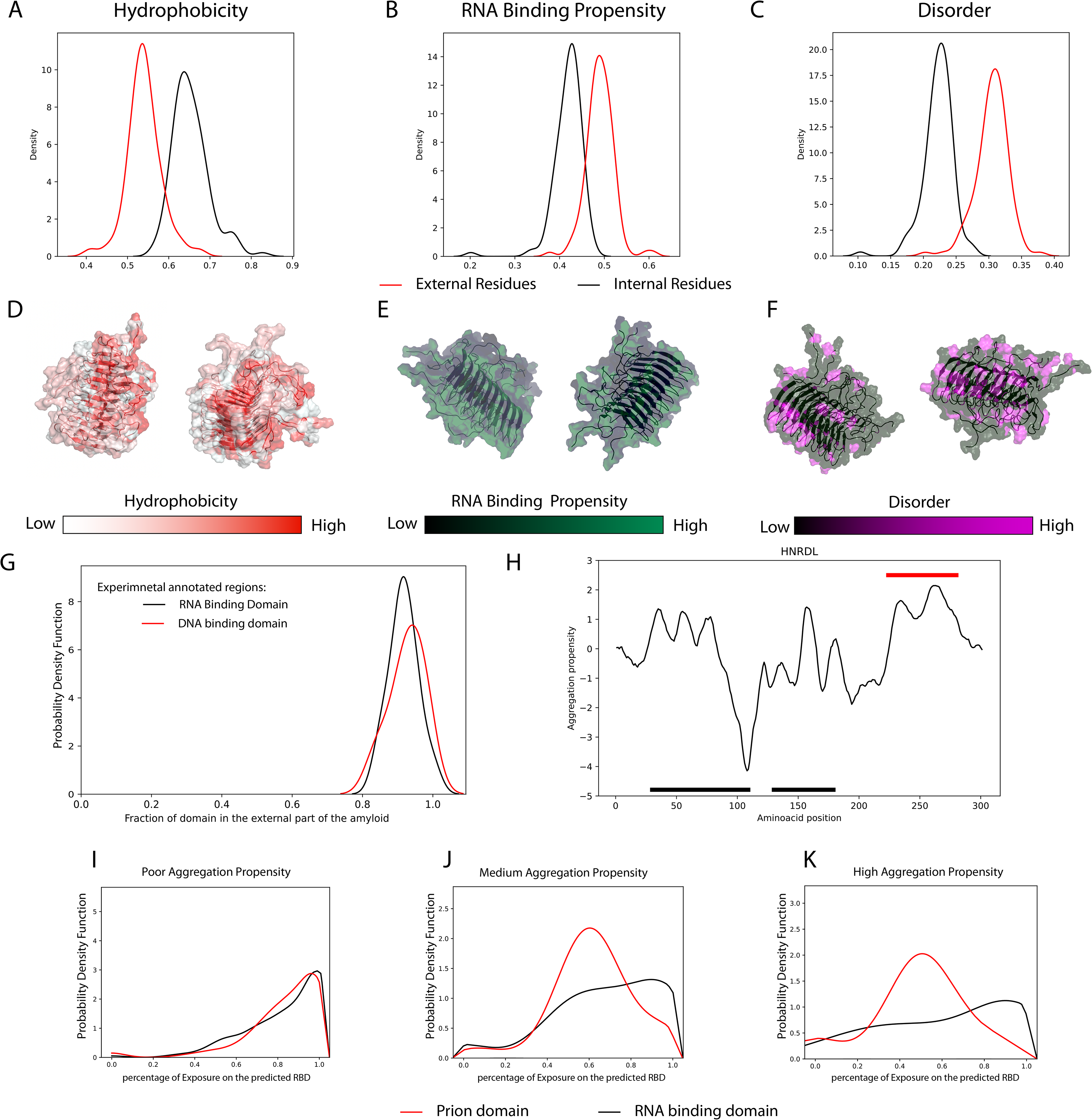
Physico-chemical properties of internal and external regions of amyloids. **A-C)** Distribution score of the amyloid dataset using the Zyggregator method for internal and external portions of the amyloids (the curves are significantly different with p-value<0.05; Kolmogorov Smirnov test).. **D-F)** Molecular structure of the HET-s prion protein (PDB 2RNM) solved by solid-state NMR, depicted by different color scales for hydrophobicity, RNA-binding propensity, and disorder, respectively. **G)** Percentage of the RNA and DNA binding domains belonging to the external part of the fibril. **H)** Zyggregator profile of hnRNPDL-2. Black lines mark the RBDs and the red line the internal residues found experimentally. **I-K)** Percentage of residues exposed for the RNA-binding and prion domains for the human proteome at different aggregation propensity scores (see also **Supplementary Table 4**).

To better visualize this trend we chose the structure of the amyloid formed by *Podospora anserina* protein HET-s (PDB 2RNM) as an example (**Fig. 2D-F**). The structure was determined using solid-state NMR and included the flexible external part of the amyloid. Using different color scales, the distribution trends are clearly visible, especially in the hydrophobic amyloid core (**Fig. 2D**). The RNA-binding propensity is considerably higher for the unstructured external portion of the fibers, similarly to the structural disorder (**Fig. 2D-F, Supplementary Fig. 2B**). Overall, the analysis of the HET-s amyloid structure shows excellent agreement with the predicted results from the computational analysis.

Within the set of amyloids (**Fig 2A-C),** we analyzed proteins that have RNA and DNA binding domains annotated in Uniprot (**Supplementary Tables 2 and 3**). Consistent with our calculations of RNA-binding abilities, the nucleic acid-binding domains are predicted to be external to the aggregate (**Fig. 2G**). This is well exemplified by hnRNPDL-2, a protein in which residues 224-280 were found by cryoEM to constitute the core of the fibril (PDB 7ZIR), while two RNA-binding domains are external^12^. The aggregation propensity predicted by Zyggregator captures the inner part of the aggregate very well, and the RNA-binding domains are indeed predicted to be more exposed (**Fig. 2H**). Further investigating this phenomenon at the human proteome level, we carried out predictions of prion and RNA-binding domains respectively using PLAAC^51^ and *cat*RAPID signature^63^. Proportionally to the overall aggregation propensity computed with Zyggregator, we observed that prion domains are in the core of the fibril while RNA-binding domains tend to be external (**Fig. 2 I-K, Supplementary Table 4**). In agreement with our analysis of hnRNPDL-2, this finding indicates that prion domains are internal to the amyloid core, while RNA-binding domains are external.

### Physico-chemical properties of alpha synuclein aggregates

We next analyzed the aggregation propensity of α-synuclein (aS or aS140), a protein with a well-characterized amyloid core^41^. We computed both the overall score of the wild type sequence (**Fig. 3A**) and performed a mutational analysis with 10 000 random single point variations^64^. In accordance with the data shown in the literature^65^, the central region of the protein is crucial for protein aggregation, with a peak coinciding with the NAC region (residues 61-95) and individual peaks corresponding to the three-repeat region, as shown by Doherty and colleagues^24^ (**Fig. 3A**).

**Figure 3.**
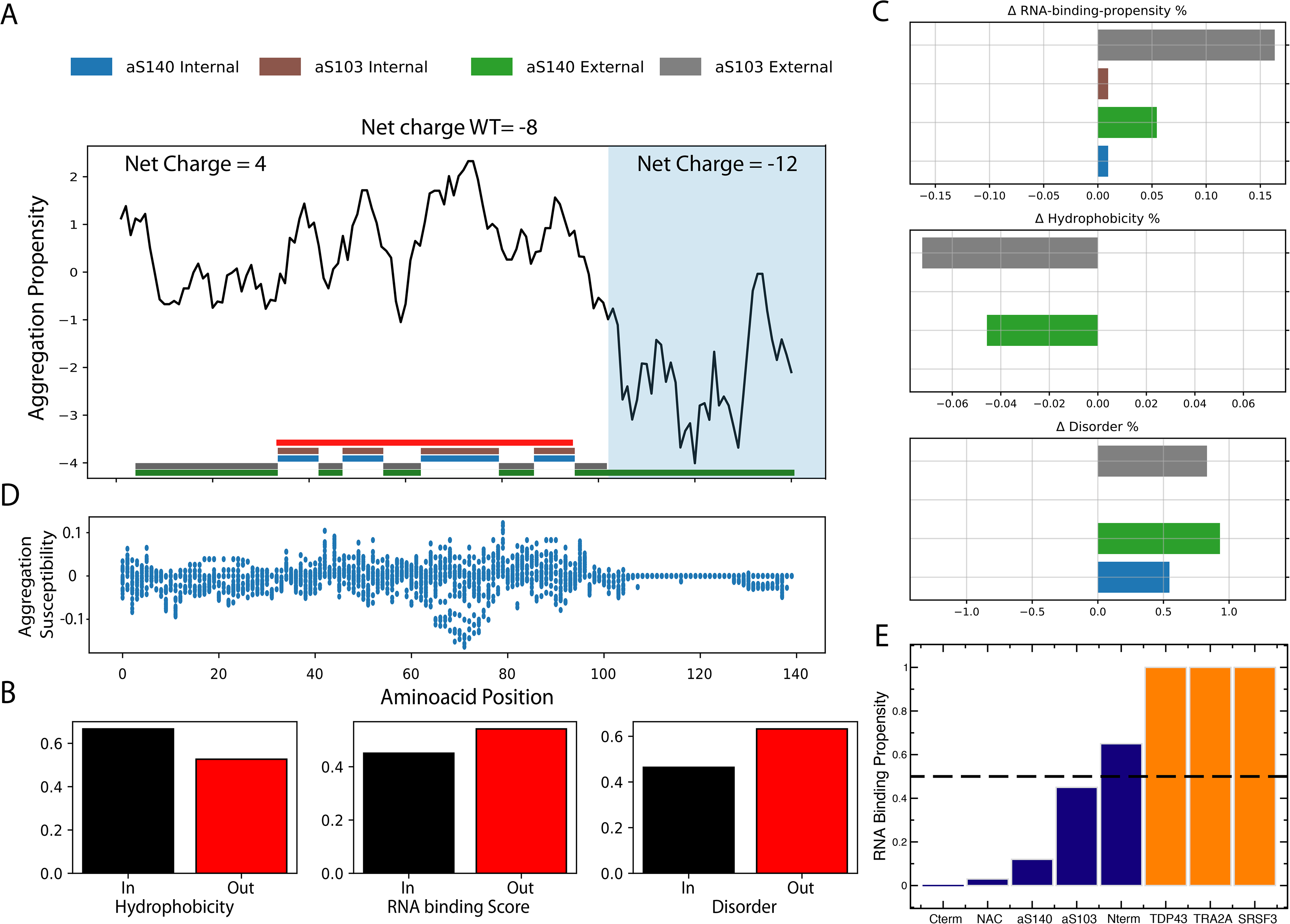
Aggregation propensity of aS140 and aS103. **A)** Zyggregator profile of aS140, with colored bars indicating predicted internal and external parts. The experimentally determined amyloid core of aS is marked with a red bar. **B)** Physico-chemical properties of aS140 and aS103. **C)** Random mutagenesis analysis shows that the NAC region and N-terminus are susceptible to significant changes in aggregation propensity. **D)** Hydrophobicity, RNA-binding propensity, and disorder of aS relative to its internal and external parts. Differences with respect to the mean of the amyloid histograms are shown (**Fig. 2A-C**). **E)** RNA-binding predictions of different regions of aS140 and aS103, compared with canonical RBP (scores > 0.5 indicate non-negligible interaction propensities).

Mutations in the NAC region display a strong impact on aS140 aggregation (**Fig. 3B**) and the external regions of aS140 aggregates are predicted to be disordered with low overall hydrophobicity and able to interact with RNA molecules (**Fig. 3C**). Amino acids 67-82 appear to be particularly sensitive to change and dramatically affect the propensity of aS to aggregate, in agreement with previous experimental work^66^. Random single mutations in the C-terminus have instead negligible effects. Yet, 14 out of the 24 negatively charged residues in aS140 (18 Glu and 6 Asp) occur in this region, indicating that the truncated form, lacking all amino acids in positions 103-140 (aS103), could behave differently compared to the wild type protein in terms of aggregation propensity as well as RNA-binding ability.

Computing the scores of the physico-chemical properties for both aS140 and aS103 reveals a strong difference between the two (**Fig. 3D**). The removal of the 12 negatively charged residues in the C-terminal portion results in a net charge increment (**Fig. 3D**). Accordingly, the external regions of aS103 and aS140 show a lower degree of hydrophobicity compared to the average of the dataset and an increase in RNA-binding ability (**Fig. 3D)** due to the abundance of charged and polar residues outside the NAC region. We note that removing the C-term slightly affects the predicted structural disorder, which is consistent with the overall unfolded nature of the whole aS.

We used the *cat*RAPID approach^56,58^ to estimate the ability of aS140 and aS103 to interact with RNA (**Materials and Methods**). While the C-terminus and NAC show low interaction propensity, the N-terminal domain is predicted to be external to the aggregate and able to weakly interact with RNA (**Fig. 3E**). Based on this observation, we hypothesize that aS might be able to sequester RNA upon aggregation. As we will show in the following sections, this effect could shape the dynamic of the aggregation propensity, tuning the concentration of the RNA in solution.

### RNA affects the aggregation of alpha synuclein isoforms in different ways

To study experimentally the effect that RNA might have on the aggregation kinetics of aS140 and aS103, we performed aggregation assays in the presence of a fluorescent aggregate intercalator. Both aS103 and aS140 were incubated with increasing concentrations of total RNA, from 0 to 500 ng/μL, for 24 h (**Fig. 4A; Supplementary Fig. 3A-B**). Since the experiments were performed in the presence of a reporter dye, visualization of the aggregates at the end of the experiments with a fluorescence microscope confirmed the presence of protein aggregates in all samples and in every tested condition (**Supplementary Fig. 4A**).

**Figure 4.**
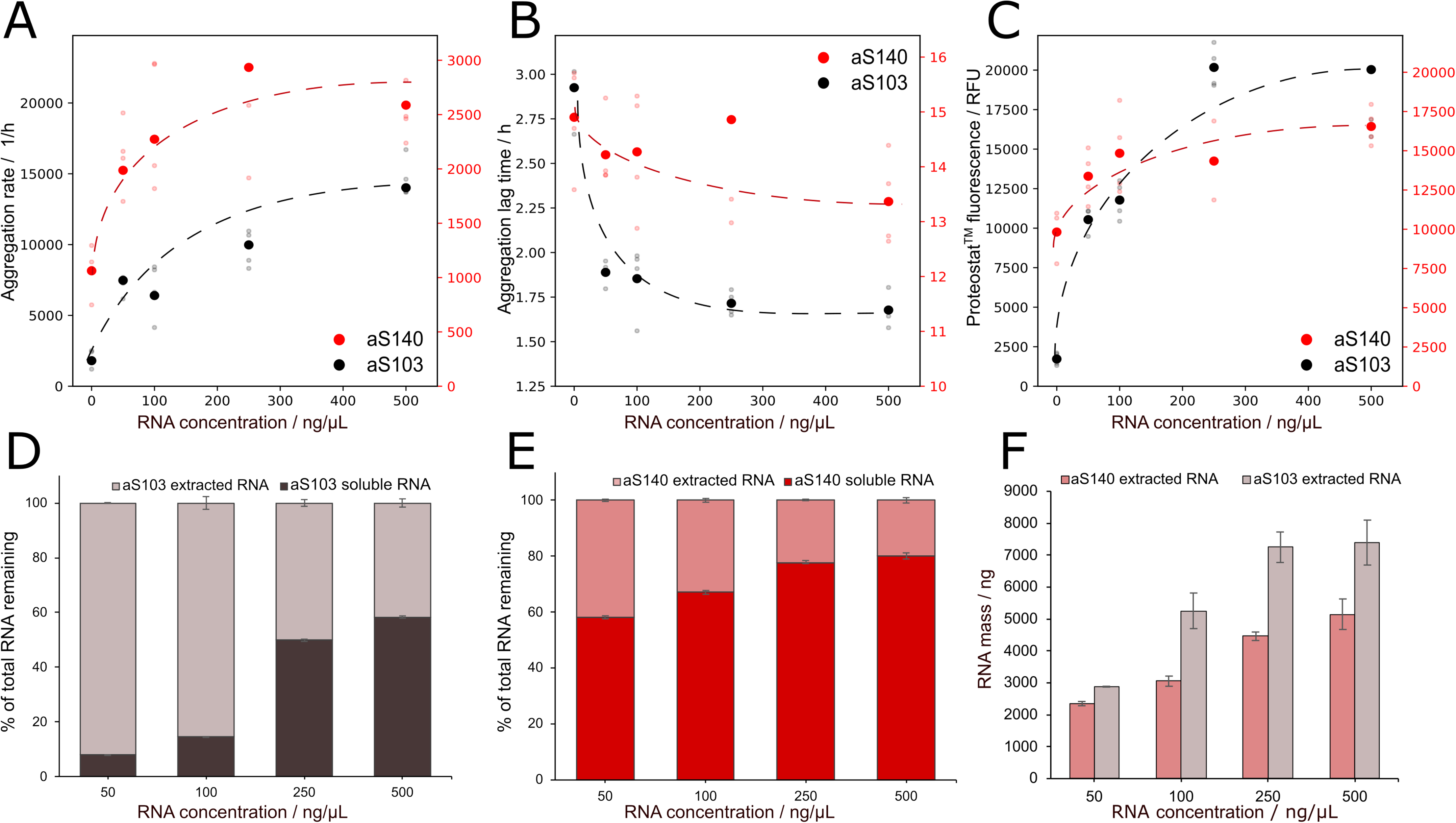
In vitro aggregation assays of aS140 and aS103 in the presence of RNA. **A**) Calculated aggregation rates from aggregation data (**Supplementary Fig. 3**) of time resolved Proteostat™ fluorescence. Data shown are calculated from parameters, derived from the fitted curve to 6 technical replicates of 3 separate experiments (see **Materials and Methods**). The presence of RNA increases the rate of aggregation for both proteins, with the aS140 rate reaching a plateau with concentrations higher than 250 ng/μL. aS103 has a significantly shorter aggregation lag time than aS140. **B**) The addition of RNA further decreases the lag phase, however increasing concentrations do not seem to drastically affect if further while the opposite is true for aS140. **C)** Increasing concentration of RNA also elevates the plateau of fluorescence for both proteins, especially significantly for aS103. **D and E)** Quantification of nucleic acids from aS aggregation assays indicates different behavior for aS140 and aS103. Comparing RNA in solution and the fraction extracted from aggregates of aS103 and aS140, we found that RNA is preferentially sequestered by aS103 aggregates. **F)** The average amount of RNA extracted from aS103 and aS140 aggregates at different initial starting concentrations show again that aS103 aggregates have a higher propensity of sequestering soluble RNA upon aggregation.

As previously reported in literature, aS103 aggregates significantly faster than aS140, irrespective of the concentration of RNA (**Fig. 3A and B**)^67^, in agreement with our predictions. In the presence of RNA, the elongation phase of aS103 is significantly faster than that of aS140 (**Fig. 4A**) and is accompanied by a consistently shorter lag phase (**Fig. 4B**). Our data analysis shows that the estimated aggregation rate of aS103 is approximately two-fold higher in the absence of RNA and up to 10-fold higher in the presence of 500 ng/μL total RNA extract. Importantly, increasing concentrations of RNA lead to varied effects on the aggregation rate of both proteins. For aS140, the aggregation rate in the presence of RNA initially scales exponentially, increasing ca. five-fold at a concentration 250 ng/μL. Then the rate seems to reach a plateau and does not increase further (**Fig. 4A**). By contrast, incremental additions of RNA monotonically increases the aggregation rate of aS103 over ten-fold, reaching a maximum at 500 ng/μL with no evidence of reaching a plateau (**Fig. 4A**). RNA affects the lag time in the opposite way, with increasing concentrations effectively decreasing the lag time of aS103 until reaching a stable plateau at 1.6 h. The opposite is true for aS140, where higher concentrations only start drastically decreasing the lag time with RNA concentrations higher than 250 ng/μL (**Fig. 4B**). The actual maximum value of fluorescence at the end of the exponential phase however does not differ much between aS103 and aS140 (**Fig. 4C)**. RNA incrementally increases the level of maximum fluorescence by additions up to 250 ng/μL when the value stabilizes. These results suggest not only that RNA has a profound effect on aS aggregation, it also affects aS140 and aS103 in different and dynamic ways as our computational analysis predicted.

Having observed the effect RNA has on the aggregation kinetics of aS, we then set to test the original hypothesis that aS can sequester the RNA upon aggregation. First we verified that the monomeric forms of both proteins do not interact with RNA (**Supplementary Fig. 4B**).

We then quantified the RNA before and after the aggregation assays in the soluble and insoluble fractions as described (**Materials and Methods**). The concentration was measured individually in each well and normalized according to the initial one. Because of differences in the volumes in different fractions (soluble and insoluble), concentrations were recalculated into RNA mass and the data presented as the percentage of total RNA mass after aggregation in different fractions (**Fig. 4D and E**). The results clearly indicate that both aS140 and aS103 are able to sequester RNA through aggregation. Furthermore, there is a clear difference between the amount of RNA sequestered by individual protein constructs. As predicted by the computational analysis, aS103 aggregates can sequester RNA to a much higher degree than aS140 (**Fig. 4F**). The difference is less noticeable at lower RNA concentrations, but becomes more pronounced at higher concentrations. As the starting concentration of the protein is always 50 μM, it is safe to assume the difference is due to the higher sequestration capacity of aS103 aggregates formed. This is further supported by the fact that less than 10 % of the starting amount of RNA is found soluble at the concentration of 50 ng/μL (**Fig. 4D**). This sequestration capacity seems to be saturated at 250 ng/μL for aS103 while it keeps increasing for aS140 (**Fig. 4F**).

To verify whether the different distribution of RNA among the soluble and insoluble fractions is specific to protein aggregation, we repeated the experiments in the same conditions with bovine serum albumin, a non-aggregating, non-RNA-binding protein. The quantification of RNA in both conditions showed comparable values to the control values of total RNA without any protein added (**Supplementary Fig. 5A**). The wells in both cases were inspected by fluorescence microscopy and no aggregates were detected (**Supplementary Fig. 4A).**

We repeated the aggregation of aS103 and aS140 in the presence of total cell DNA extract, to determine whether what we observed is an RNA-specific effect. Since DNA and RNA degrade at different rates, we adjusted the normalization to account for it (**Supplementary Fig. 5B**). aS103 still shows the ability to sequester DNA at lower initial DNA concentrations, however higher concentrations apparently limit the binding ability (**Supplementary Fig. 5C**). aS140 barely sequesters any DNA in the aggregates, with less than 10% extracted from the insoluble part (**Supplementary Fig. 5D).**

These results show that both aS140 and aS103 have the ability to sequester nucleic acids upon aggregation. The grade of incorporation is protein sequence-dependent and confirms the computational analysis predicting aS103 aggregates as the higher-propensity binders. The results further emphasize that protein aggregation is a dynamic process, and that aggregated species not only display different biophysical features but could also acquire new functional properties distinct from the ones attributed to the monomers.

## DISCUSSION

Aggregates are able to sequester other molecules, including proteins^1^ and several metabolites^68^. Here, we tested the hypothesis that RNA could be trapped by amyloid aggregates. Our hypothesis was formulated considering that aggregation-prone regions of proteins are mostly hydrophobic and buried in the core of fibrils^41^, while external regions, prevalently hydrophilic and structurally disordered, can interact with nucleic acids, if their electrostatics are favorable^69^. Several lines of evidence, both computational and experimental, seemed to confirm our idea. Regions outside the core of amyloids are predicted to interact with RNA and annotated RNA-binding domains have a strong propensity to be external with respect to the amyloid core. The recently reported cryoEM structures of tau and hnRNPDL-2 show that RNA can be trapped on the surface of the aggregate^13^ and the RNA-binding domains are outside the amyloid core^12^.

Intriguingly, a number of artificial nucleic acid molecules, RNA and DNA aptamers, have been designed to impact protein aggregation^4,70^. For instance, an aptamer called T-SO517, developed by Ikebukuro and colleagues against oligomeric αS140, bound not only to the target but also to Aβ40 oligomers^71^. This finding confirms our hypothesis that aggregates can bind to nucleic acids, although it indicates that the binding may be non-specific. Thus, it is possible to speculate that sequestration of RNA material in the cellular context could contribute to the toxicity of amyloid aggregates, a hypothesis that has been previously formulated for proteins^1^.

Here, we reasoned that the formation of the amyloid core of aS fibrils, involving the most hydrophobic residues (i.e., the NAC region), leaves the charged amino acids (i.e., N-terminal and C-terminal) available for interactions with other molecules such as RNA. This is corroborated by a number of experiments such as site-directed spin labeling coupled with EPR, hydrogen/deuterium exchange, and limited proteolysis^34,48–50^, indicating that the central part of the protein is engaged in forming the core of the aggregates while the rest is exposed to the solvent. We predicted that removal of the negative charges corresponding to amino acids 104-140, not included in the core of the aggregate, would sharply increase the RNA-binding propensity of the regions outside the NAC. Thus, we tested the hypothesis of RNA-binding gain-of-function upon aggregation.

Our experimental results confirm the computational observation of acquired RNA sequestration ability and identify it as a way of modulating aS aggregation process by directly affecting the aggregation rates. Sequestration is protein-sequence dependent: aS103 shows a higher sequestration propensity compared to wild type aS140, likely due to the increased electrostatic attraction. RNA also modifies the aggregation rate of both aS variants.

As previously reported, the key to modulating aS aggregation propensity lies in the long-range intra and intermolecular interactions between the N-terminal and C-terminal domains of the protein, which shield the NAC region and prevent its self-assembly^34,35^. RNA likely enters in contact with the N-terminal, thus interfering with these interactions. The truncated aS103 lacks the C-terminal domain and *a priori* cannot engage in the shielding effect of the NAC region. It is plausible that, through transient interactions via the N-terminal domain, RNA acts as a molecular bridge between protein molecules, bringing the NAC regions into closer contacts. As C-terminal truncations of aS are commonly found in brain samples from patients and aS103 is prevalent in the medial temporal lobe in Lewy body dementia, we speculate that RNA might play a role in the onset of the disease^28^.

Very recently, the combined solid-state NMR - cryoEM approach to study the fibrils of aS by Zhang and colleagues has confirmed our observations regarding the outer disorder of aS fibrils ^72^. The structure (PDB 7YK2) however also highlighted the ordering and interaction between the core of the fibril and the G14 - G25 region in the N-terminal part. This suggests that additional ordering of the structure might be present in the N-terminus upon aggregation. It is important to note, however, that the fibrils in the study were grown in the absence of RNA, which could potentially prevent these additional ordering to occur upon misfolding by shielding the N-terminus.

We note that RNA concentrations in our experiments are near-to-physiological. Indeed, we considered a mammalian cell has a cell volume between 1,000 and 10,000 μm^3^ (i.e., 10^-6^ μL −10^-5^ μL ^73^) and an RNA amount of 10 - 30 pg (i.e., 0.01 - 0.03 ng ^74^), which indicates that 500 ng/μL of RNA are in the expected normal range (i.e., 0.01 ng in 10^-5^ μL, which is 1000 ng/μL). Monomeric, physiological aS was found associated with nucleic acids^20,75^, nucleosomes^76^ and processing bodies^77^. However, no proof of direct and functional RNA binding by the cytosolic, aggregated aS has ever been discovered. While Siegert and colleagues have shown RNA has little effect and can actually abrogate aS phase separation^78^, the molecular aspects of the effect of RNA on aS liquid-to-solid phase transition was not characterized. It should be mentioned that Cohlberg and colleagues have previously reported that negatively charged molecules such as heparin display a stronger effect on aS aggregation compared to 1μg/μL RNA^38^. Nevertheless, it must be noted that the authors used thioflavin T to quantify aS aggregation. It has been reported that this dye can interact also with RNA^79^, interfering with the correct quantification of protein aggregation. For this reason, we selected a different aggregation reporter, Proteostat™, that binds to RNA only weakly^80^, obtaining significantly different results. An intriguing aspect is the pathological implication of RNA sequestration in synucleinopathies.

Overall our study shows that aggregation and the misfolding associated with it can lead to different biophysical and functional properties compared to the monomeric soluble protein forms. aS, a non-RNA-binding protein, can acquire the ability to sequester RNA in a sequence-dependent manner, bringing further emphasis on the dynamic structural and functional transitions between different protein states.

## ACKNOWLEDGMENTS

The authors would like to thank the ‘RNA initiative’ at IIT and all the members of Tartaglia’s lab at CRG, Sapienza and IIT. Our research was supported by the ERC ASTRA_855923 (GGT) and H2020 projects IASIS_727658 and INFORE_825080 (GGT). EZ received funding from the Newton fellowship scheme and the MINDED fellowship of the European Union’s Horizon 2020 research and innovation program under the Marie Skłodowska-Curie grant agreement No. 754490.

## MATERIALS AND METHODS

### Amyloid Dataset

Amyloid proteins (**Supplementary Table 1)** were taken from the PRALINE database^40^. The internal and external parts of amyloids were computed using Zyggregator^41^. In **Fig. 1**, we report the comparison of predicted and experimentally validated internal and external regions of calcitonin^42,43^, amylin^44^, glucagon^45^, Aβ42^46,47^, α-synuclein^48–50^, hnRNPDL-2^12^ and tau^13^. The optimal threshold is in the range 0.5-1.5 and corresponds to an Area under the Roc Curve of 0.84.

### Analysis of the inner and outer part of amyloids

For each residue in a given sequence (**Supplementary Table 1)**, we used the Zyggregator that predicts the individual contribution to aggregation based on predictions of hydrophobicity, secondary structure and ^41^. Our previous work indicates that the residues with high aggregation propensities are buried in the fibril, while those with low aggregation propensity are exposed to the solvent^41^.

### RNA/DNA binding domains and aggregation

We exploited annotations of RNA/DNA binding domains available from UniProt to characterize their physico-chemical properties in relation to the aggregation propensity (**Supplementary Tables 2 and 3, Fig. 2**). Specifically, we computed the fraction of RNA/DNA domains belonging to the predicted external part of the amyloids. The same analysis was carried out for prion domains predicted using PLAAC^51^, as in a previous work^9^ (**Fig. 2**).

### Random mutagenesis

We performed an *in silico* random mutation analysis to define the role of each residue of aS sequence in the aggregation propensity of the protein. We selected random positions in the sequence and replaced them, one by one, with amino acids from an uniform distribution. We used the Zyggregator algorithm^41^ to compute the aggregation propensity of the protein before and after the mutation since, for each residue in a given sequence, the method predicts the individual contribution to aggregation^41^. We simulated 10000 random mutations and calculated for each one the difference with the wild type in terms of Zyggregator score, expressed as a percentage.

### Physico-chemical properties of external regions

Calculations of hydrophobicity^52^, disorder^53^ and RNA-binding ability^54^ have been carried out through the CleverMachine approach that predicts physico-chemical properties of protein sequences^55^.

In the Supporting Information (**Supplementary Fig. 1 and 2**), we show that the particular scale used is not strictly important, as the behavior of the histograms of the chemical physical property did not change with the scale change.

### *cat*RAPID predictions of protein-RNA interactions

In *cat*RAPID, the interaction propensity between a protein-RNA pair is computed using secondary structure properties, van der Waals and hydrogen bonding potentials^56^. The algorithm is able to separate interacting versus non-interacting pairs with an area under the curve (AUC) receiver operating characteristic (ROC) curve of 0.78 (with false discovery rate (FDR) significantly below 0.25 when the Z-score values are >2)^57^. In this work, the N-terminus of aS (residues 1-25) is predicted to attract RNA, while the C-terminal region (residues 104-140) shows much poorer propensity. In our calculations^58^, we used the human transcriptome and computed the overall interaction propensity (>0.4 corresponding to low binding affinity) for different aS regions. TDP43, TRA2A and SRSF3 are used as a positive control. aS103 and aS140 are not predicted to interact as monomers.

### Protein production

The cDNA for the aS140 was inserted in a pET21a vector under the control of the T7 promoter. The aS103 cDNA was produced by deletion of the appropriate sequence from the full length construct with specific PCR primers (aS103for: 5’-TAAGCGGCCGCACTCGAGCAC-3’ and aS103rev: 5’-ATTCTTGCCCAACTGGTCCTTTTTGACAAAGC-3’).

Competent *E.coli* BL21 [DE3] cells were transformed with both constructs. 2 L of fresh LB broth supplemented with 50 μg/mL ampicillin were inoculated with overnight cultures with a 100:1 ratio. Cultures were grown at 37 °C up to an optical density of ca. 0.7 at 600 nm. Protein expression was induced with 0.5 mM IPTG and the cultures were shaken for 4 hours. Cells were harvested by centrifugation at 4500 g at 4 °C for 30 min, afterwards the pellet was resuspended on ice in 20 mM potassium phosphate buffer pH 7.2 and frozen immediately.

For protein purification the cells were thawed in ice-cold water and lysed by boiling at 95 °C for 30 min. The boiled suspension was centrifuged at 18000 g at 4 °C for 30 min, then streptomycin sulfate was added to the supernatant up to a concentration of 10 mg/mL. The solution was centrifuged at 18000 g at 4 °C for 30 min and the pellet containing precipitated impurities was discarded. The protein was slowly precipitated by addition of crystalline ammonium sulfate up to 360 mg/mL and the suspension was centrifuged again at 18000 g at 4 °C for 30 min. The pellet was resuspended in 20 mM Tris-HCl pH 7.4 with 400 mM KCl and dialysed against 20 mM Tris-HCl pH 7.4 overnight at 4 °C.

The proteins were purified using heparin affinity chromatography on a HiTrap™ Heparin column to remove remaining nucleic acids. Proteins were eluted with heparin elution buffer (20 mM potassium phosphate pH 7.2, 800 M NaCl) and loaded onto a HiLoad^TM^ 16/600 Superdex™ 75 gel filtration column equilibrated in 20 mM potassium phosphate pH 7.2, 100 mM KCl. The eluted proteins were pooled and checked for concentration with BCA assay and purity with spectropolarimetry and SDS PAGE, before being stored aliquoted at - 80 °C.

### Protein aggregation assays

A frozen protein aliquote of aS140 or aS103 was quickly thawed and filtered with a 0.22 μm syringe filter immediately before each assay. Total yeast RNA (totRNA, Roche) was resuspended in aggregation assay buffer (20 mM potassium phosphate pH 7.2, 100 mM KCl, 5 mM MgCl_2_). For aggregation assays in the presence of DNA, total DNA from herring sperm (Sigma Merck) was resuspended in sterile, nuclease-free water for the stock solution. Protein, totRNA and DNA were diluted to the final concentration in the aggregation assay buffer with a 1000-time diluted Proteostat™ detection dye. For the BSA control, ultra-pure, DNA- and RNA-free BSA was used (Sigma Merck) and diluted to the final concentration in the sample buffer as described above for aS. The samples were distributed in 6 replicates on a 96-well black plate with transparent bottom and a single borosilicate glass bead was added to each well to ensure sample homogeneity and reproducibility. The excitation wavelength was set to 505 nm and the emission to 590 nm. Plates were incubated at 37 °C with constant double-orbital shaking at 200 rpm. Reads were taken every 10 min for aS140 and every 5 min for aS103 for 24 hours, with each read represented as an average of 10 scans.

All protein samples were aggregated in parallel to total RNA alone at the same starting concentrations and conditions for RNA degradation control and the values reported have already been accounted for degradation.

### Aggregation data analysis

Data from the aggregation assays, obtained with Proteostat™ fluorescence dye, were fitted into a sigmoid curve using the Hill function (1):

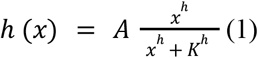

where A, h and K are the fitting parameters.

To compute the aggregation rate, we determined the slope of the fitted curves before plateau. With the previously set parameters and a Taylor expansion of *h*(*x*) around *K*, we calculated the slope of the aggregation α curve as (2):

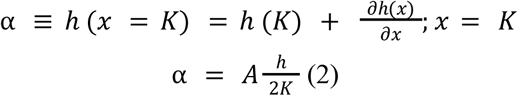

We define the “delay time” as the intersection of the fitting line with slope α centered in *x* = *K* and 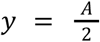. The delay time τ is defined resolving the linear system (3):

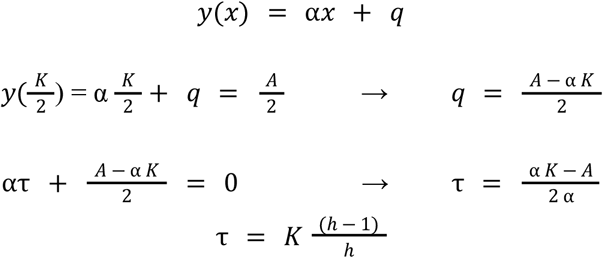

### Nucleic acid quantification and extraction from protein aggregates

After aggregation, individual replicates were transferred from the 96-well plate to Eppendorf tubes and centrifuged at 4 °C, 18.000 g for 30 min. The soluble fraction was transferred to another tube, RNA found in solution was quantified using Qubit RNA BR, and DNA with Qubit DNA BR kit (Thermo Fisher Scientific, USA).

For RNA extraction, the insoluble pellet was resuspended in 25 μL of digestion buffer (2 M urea, 100 μg/mL proteinase K, 3 mM DTT) and incubated at room temperature for 15 min. 125 μL of 6 M guanidine hydrochloride was added to a final concentration of 5 M and the sample was incubated at room temperature for further 30 min. 1 mL of Trizol™ was added and RNA was extracted using standard protocol by the manufacturer.

DNA was extracted from the insoluble protein pellet with the QIAamp DNA Mini Kit (Qiagen, Germany) according to the manufacturer protocol for DNA extraction from blood. The final elution was performed in 50 μL of buffer AE. Extracted nucleic acids were again quantified using the Qubit fluorescent dyes.

