## Supplementary Materials for "A Computational Approach Reveals the Ability of Amyloids to Sequester RNA: the Alpha Synuclein Case"

### SUPPLEMENTARY TABLES

**Supplementary Table 1.** Amyloid dataset with reported the internal and external residues, and the predicted aggregation rate

**Supplementary Table 2.** RNA- binding residues found reported experimentally in uniprot for the Amyloid proteins

**Supplementary Table 3.** DNA-binding residues found reported experimentally in uniprot for the Amyloid proteins

**Supplementary Table 4.** Positions of the RNA binding domain and prion domains from Gotor *et al* [1].

### SUPPLEMENTARY METHODS

#### Bio-layer interferometry measurements

The binding of the monomeric protein was assessed using bio-layer interferometry (BLI). The protein was biotinylated using the EZ-Link™ NHS-PEG4-Biotin reagent (Thermo Fisher Scientific) according to the instructions from the manufacturer. Binding experiments were performed on an Octet K2 (Sartorius) instrument. Streptavidin (SA) biosensors (Sartorius) were equilibrated in the binding buffer (20 mM K phosphate pH 7.2, 100 mM KCl, 5 mM $MgCl_2$) for 80 s. 1 μM of stock solution of bt-aS140 and bt-aS103 were prepared and loaded onto Streptavidin (SA) biosensors (300 s). After the loading, the biosensors were washed in fresh binding buffer for 300 s. Association and dissociation phases lasted for 600 s each and were performed in 500 ng/μL solution of total RNA from the same stock used for aggregation and fresh binding buffer respectively.

#### Aggregate microscopy

Imaging of the aggregated species was performed on a Nikon Eclipse Ts2 inverted microscope directly from the wells of the plate where aggregation was performed. Images were acquired at 20x magnification, with 200 ms exposure at 560 nm laser excitation for Proteostat™ dye. 3 images were acquired per well and the most representative chosen.

### SUPPLEMENTARY FIGURES

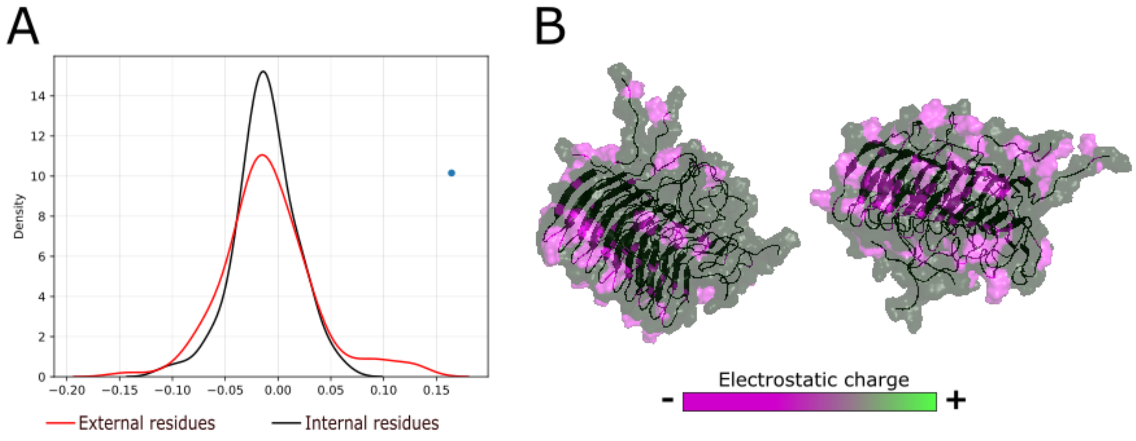


**Supplementary Figure 1.** *Analysis of amyloid dataset according to electrostatic charge.* **A**) Distribution of the net charge ofaminoacid residues of the amyloid dataset shows an overlap and no significant difference between the internal (black) and external (red) parts. **B**) The structure of the amyloid of *Podospora anserina* protein HET-s (PDB 2RNM) shows that charged residues are outside the core of the aggregate.

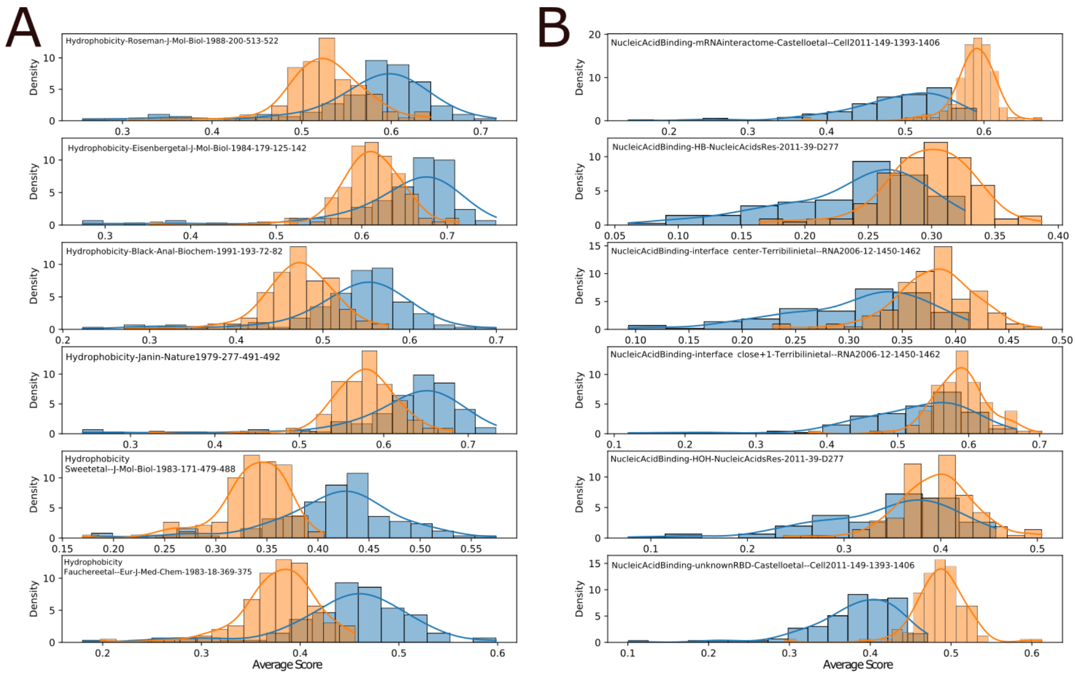


**Supplementary Figure 2.** *Analysis of amyloid dataset according to hydrophobicity and nucleic acid binding propensity for the external (orange) and internal (blue) residues.* **A)** Comparison of the CleverMachine predictors indicates the external residues of amyloids as predicted by Zyggregator have a much lower overall hydrophobicity score [2]. **B)** The opposite trend is seen for nucleic acid binding propensity, estimated always using the CleverMachine [2].

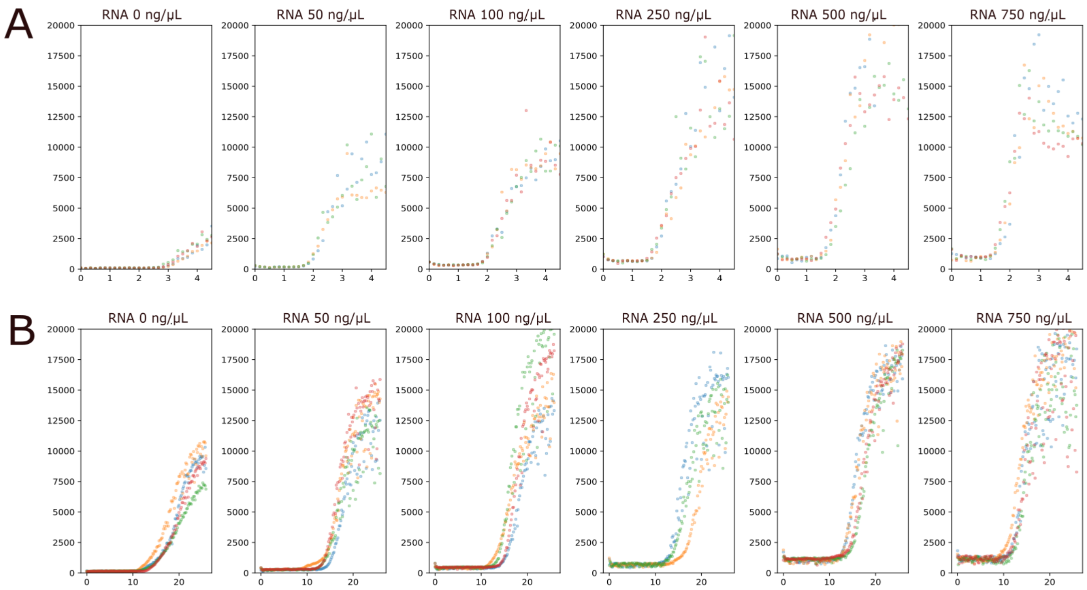


**Supplementary Figure 3.** *Raw data for aS in vitro aggregation assays in the presence of total yeast RNA with Proteostat™ dye fluorescence as the y-axis values.* **A**) aS103 shows a much faster onset of the exponential phase of growth 2 h after the start of the assay which progressively shortens with increasing RNA concentration. The plateau fluorescence for the protein without RNA is also comparatively low with a visible increase when RNA concentration is increased. **B**) An identical trend is observed for aS140. The protein without RNA has a higher fluorescence value at plateau compared to aS103, while values at higher RNA concentrations are comparable.

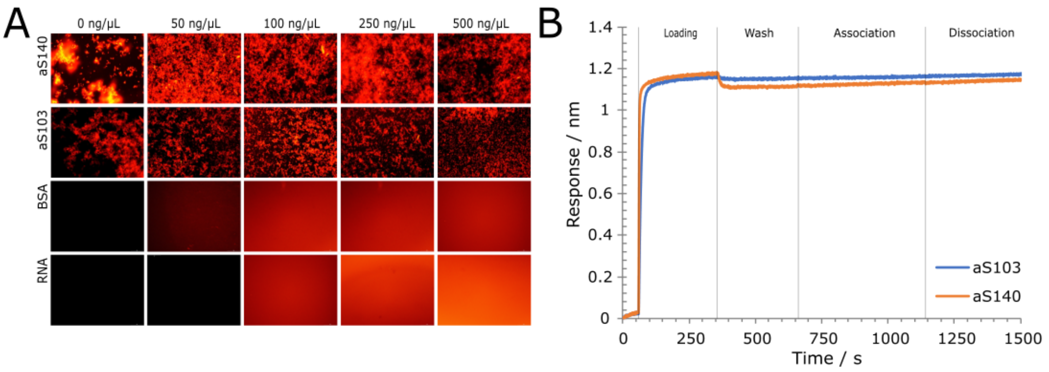


**Supplementary Figure 4.** *Control fluorescent imaging of aggregation assays and monomer-RNA interaction.* **A**) Fluorescence imaging of the wells after aggregation. Wells containing aS103 and aS140 contained visible aggregates after incubation at all concentrations. BSA and RNA incubation alone did not result in any Proteostat™-positive inclusions visible. **B**) Bio-layer interferometry graphs showing response curves for biotinylated aS103 and aS140 in the presence of total yeast RNA (500 ng/µL in 20 mM potassium phosphate pH 7.2, 100 mM KCl, 5 mM $MgCl_2$). No shift in the response signal is observed, showing the interactions between the monomeric proteins and RNA, if occurring at all, are likely to be transient and weak.

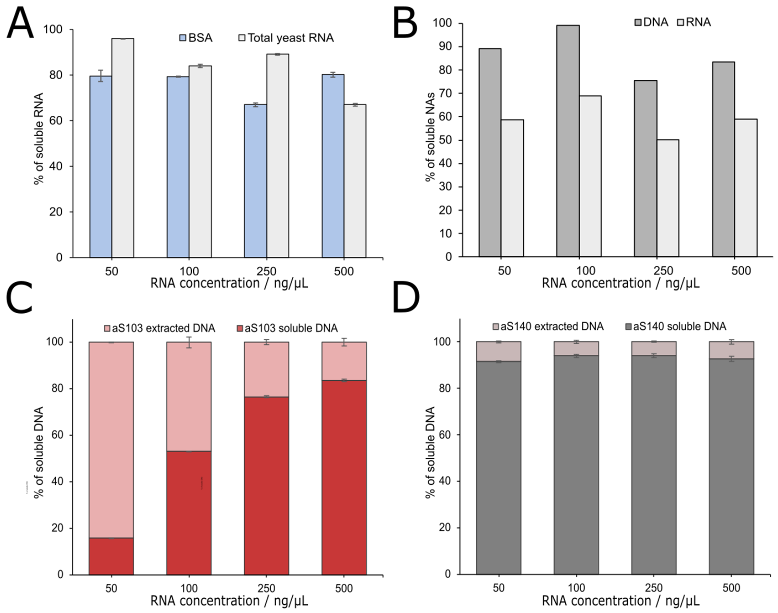


**Supplementary Figure 5.** *Quantification of soluble nucleic acids after in vitro aggregation.* **A)** Experiments with 50 µM BSA and total RNA alone show that the RNA remains mostly intact and in solution. **B)** DNA by itself is more stable in the conditions of the aggregation assays, with more than 70 % recovede at the end of the experiment. RNA gets degraded faster, and only 50 % remains after the 24-hour incubation. **C,D)** Normalized quantification of DNA fractions in aS103 and aS140 aggregates shows different trends for aS103 and aS140. **C)** aS103 aggregates sequester DNA at much higher rates compared to aS140. The capacity seems to be diminishing at higher DNA concentrations, however still more than 20 % of total DNA is sequestered at 500 ng/µL compared to less than 10 % of total DNA being extracted from the aS140 aggregates.
